# Deep learning: Using machine learning to study biological vision

**DOI:** 10.1101/178152

**Authors:** Najib J. Majaj, Denis G. Pelli

## Abstract

Today many vision-science presentations employ machine learning, especially the version called “deep learning”. Many neuroscientists use machine learning to decode neural responses. Many perception scientists try to understand how living organisms recognize objects. To them, deep neural networks offer benchmark accuracies for recognition of learned stimuli. Originally machine learning was inspired by the brain. Today, machine learning is used as a statistical tool to decode brain activity. Tomorrow, deep neural networks might become our best model of brain function. This brief overview of the use of machine learning in biological vision touches on its strengths, weaknesses, milestones, controversies, and current directions. Here, we hope to help vision scientists assess what role machine learning should play in their research.

## INTRODUCTION

What does machine learning offer to biological-vision scientists? Machine learning was developed as a tool for automated classification, optimized for accuracy. Machine learning is used in a broad range of applications (Brynjolfsson, 2018), e.g. regression in stock market forecasting and reinforcement learning to play chess, but here we focus on classification. Physiologists use it to identify stimuli based on neural activity. To study perception, physiologists measure neural activity and psychophysicists measure overt responses, like pressing a button. Physiologists and psychophysicists are starting to consider deep learning as a model for object recognition by human and nonhuman primates (Cadieu et al., 2014; Ziskind et al., 2014; Yamins et al., 2014; Khaligh-Razavi & Kriegeskorte, 2014; Testolin, Stoianov, & Zorzi, 2017). We suppose that most of our readers have heard of machine learning but are wondering whether it would be useful in their own research. We begin by describing some of its pluses and minuses.

## PLUSES: WHAT IT’S GOOD FOR

At the very least, machine learning is a powerful tool for interpreting biological data. A particular form of machine learning, *deep learning*, is very popular (Fig. 1). Is it just a fad? For computer vision, the old paradigm was: feature detection, followed by segmentation, and then grouping (Marr, 1982). With machine learning tools, the new paradigm is to just define the task and provide a set of labeled examples, and the algorithm builds the classifier. (This is “supervised” learning; we discuss unsupervised learning below.)

**Figure 1.**
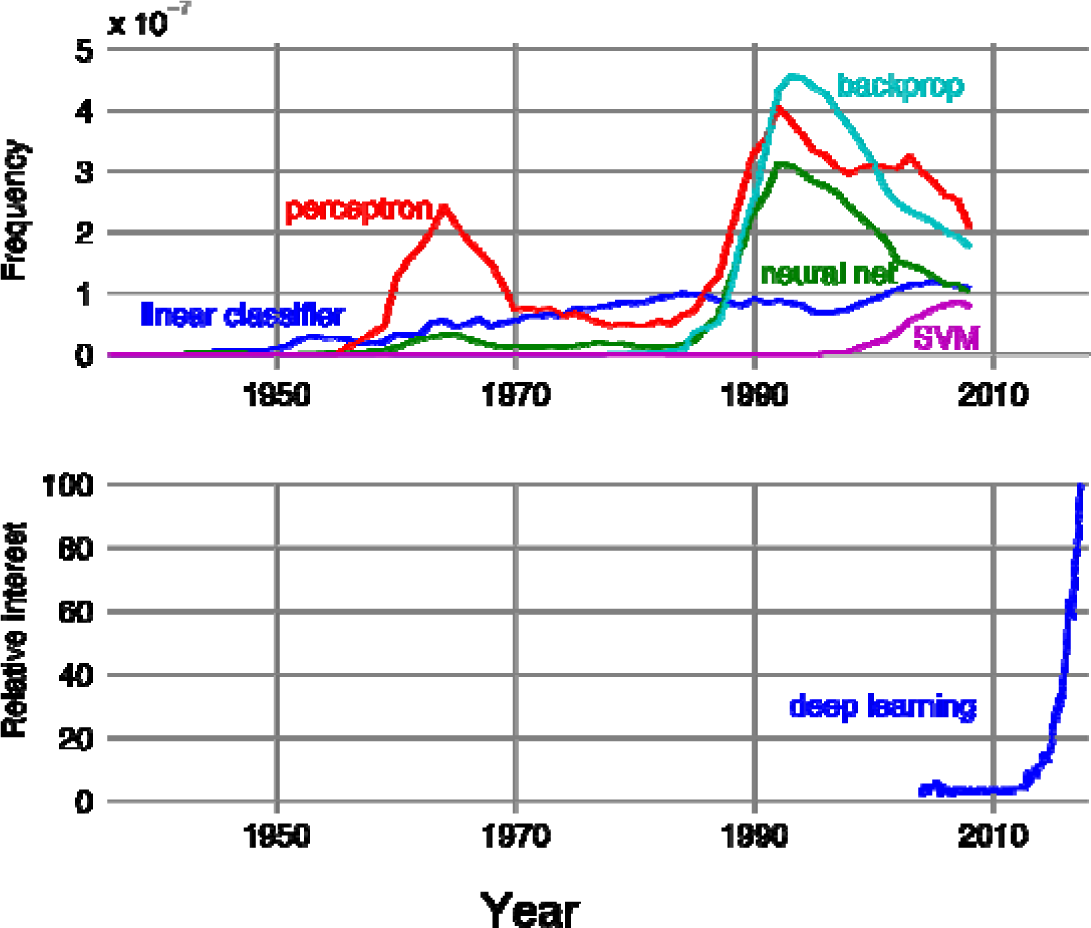
**Top:** The frequency of appearance of each of five terms — “linear classifier”, “perceptron”, “support vector machine”, “neural net” and “backprop” — in books indexed by Google in each year of publication. Google counts instances of words and phrases of *n* words, and calls each an “ngram”. Frequency is reported as a fraction of all instances of ngrams of that length, normalized by the number of books published that year (ngram / year / books published). The figure was created using Google’s ngram viewer (https://books.google.com/ngrams), which contains a yearly count of ngrams found in sources printed between 1500 and 2008. **Bottom:** Numbers represent worldwide search interest relative to the highest point on the chart for the given year for the term “deep learning” (as reported by https://trends.google.com/trends/).

**Figure 2.**
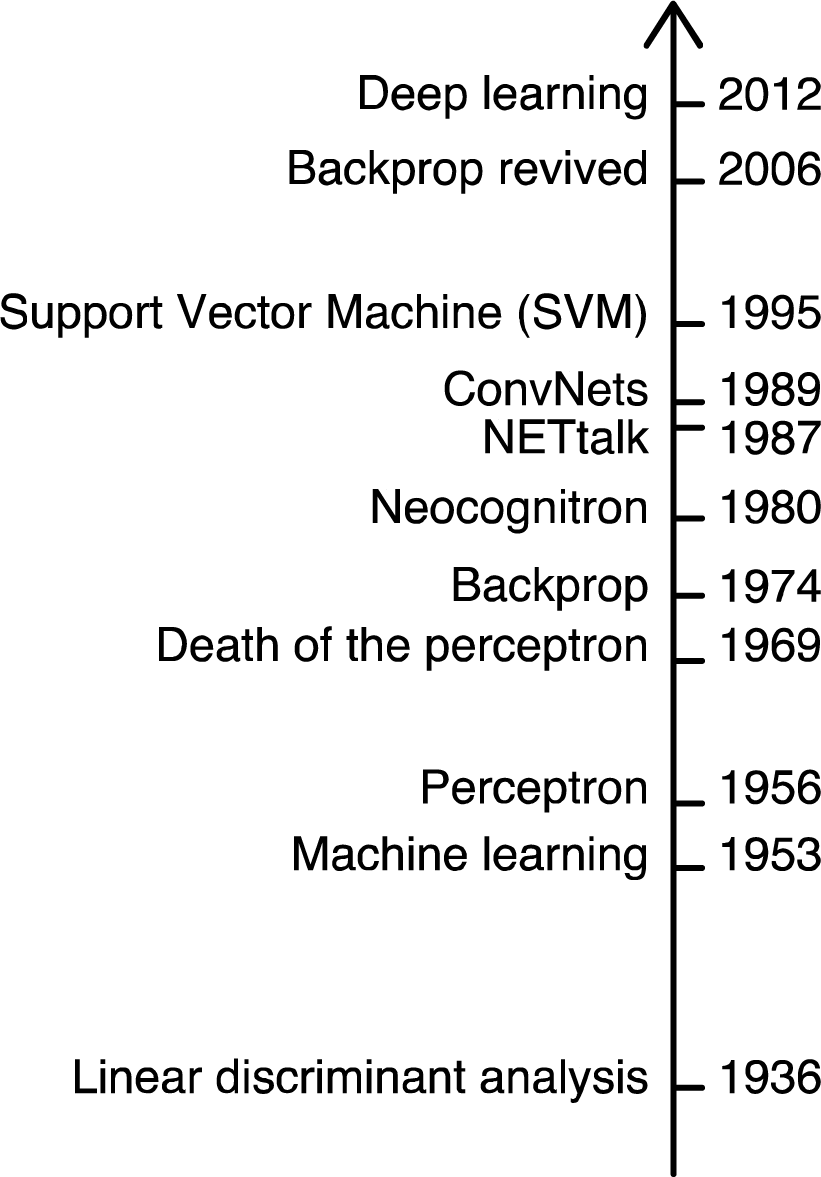
Milestones in classification.

Unlike the handcrafted pattern recognition (including segmentation and grouping) popular in the 70’s and 80’s, deep learning algorithms are generic, with little domain-specificity. ^1^ They replace hand-engineered feature detectors with filters that can be learned from the data. Advances in the mid 90’s in machine learning made it useful for practical classification, e.g. handwriting recognition (LeCun et al., 1989; Vapnik, 1999).

Machine learning allows a neurophysiologist to decode neural activity without knowing the receptive fields (Seung & Sompolinsky,1993; Hung et al., 2005). Machine learning is a big step in the shifting emphasis in neuroscience from *how* the cells encode to *what* they encode, i.e. what that code tells us about the stimulus (Barlow, 1953; Geisler, 1989). Mapping a receptive field is the foundation of neuroscience (beginning with Weber’s 1834/1996 mapping of tactile “sensory circles”). This once required single-cell recording, looking for minutes or hours at how one cell responds to each of perhaps a hundred different stimuli. Today it is clear that characterization of a single neuron’s receptive field, which was invaluable in the retina and V1, fails to characterize how higher visual areas encode the stimulus. Machine learning techniques reveal “how neuronal responses can best be used (combined) to inform perceptual decision-making” (Graf, Kohn, Jazayeri, & Movshon, 2010). The simplicity of the machine decoding can be a virtue as it allows us to discover what can be easily read-out (e.g. by a single downstream neuron) (Hung et al. 2005). Achieving psychophysical levels of performance in decoding a stimulus object’s identity and location from the neural response shows that the measured neural performance has all the information needed for the subject to do the task (Majaj et al. 2015; Hong et al. 2016).

For psychophysics, Signal Detection Theory (SDT) proved that the optimal classifier for a known signal in white noise is a template matcher (Peterson, Birdsall, & Fox, 1954; Tanner & Birdsall, 1958). Of course, SDT solves only a simple version of the general problem of object recognition. The simple version is for known signals, whereas the general problem includes variation in viewing conditions and diverse objects within a category (e.g. a chair can be any object that affords sitting). SDT introduces the very useful idea of a mathematically defined ideal observer, providing a reference for human performance (e.g. Geisler, 1989; Pelli et al., 2006). However, one drawback is that it doesn’t incorporate learning. Deep learning, on the other hand, provides a pretty good observer that learns, which may inform studies of human learning.^2^

These networks might reveal the constraints imposed by the training set on learning. Further, unlike SDT, deep neural networks cope with the complexity of real tasks. It can be hard to tell whether behavioral performance is limited by the set of stimuli, their neural representation, or the observer’s decision process (Majaj et al. 2015). Implications for classification performance are not readily apparent from direct inspection of families of stimuli and their neural responses. SDT specifies optimal performance for classification of known signals but does not tell us how to generalize beyond a training set. Machine learning does.

## MINUSES: COMMON COMPLAINTS

Some biologists point out that neural nets do not match what we know about neurons (e.g., Crick, 1989; Rubinov, 2015). Biological brains learn on the job, while neural networks need to converge before they can be used. Furthermore, once trained, deep networks generally compute in a feed-forward manner while there are major recurrent circuits in the cortex. But this may simply reflect the different ways that we use artificial and real neurons. The artificial networks are trained for a fixed task, whereas our visual brain must cope with a changing environment and task demands, so it never outgrows the need for the capacity to learn.

It is not clear, given what we know about neurons and neural plasticity, whether a backprop network can be implemented using biologically plausible circuits (but see Mazzoni et al., 1991, and Bengio et al., 2015). However, there are several promising efforts to implement more biological plausible learning rules, e.g. spike-timing-dependent plasticity (Mazzoni et al., 1991; Bengio et al., 2015; Sacramento, Costa, Bengio, & Senn, 2017).

Engineers and computer scientists, while inspired by biology, focus on developing machine learning tools that solve practical problems. Thus, models based on these tools often do not incorporate known constraints imposed by biological measurements. To this, one might counter that every biological model is an abstraction and can be useful even while failing to capture all the details of the living organism.

Some biological modelers complain that neural nets have alarmingly many parameters. Deep neural networks continue to be opaque. Before neural-network modeling, a model was simpler than the data it explained. Deep neural nets are typically as complex as the data, and the solutions are hard to visualize (but see Zeiler & Fergus, 2013). However, while the training sets and learned weights are long lists, the generative rules for the network (the computer programs) are short. Traditionally, having very many parameters has often led to overfitting, i.e. good performance on the training set and poor performance beyond it, but the breakthrough is that deep-learning networks with a huge number of parameters nevertheless generalize well. Furthermore, Bayesian nonparametric models offer a disciplined approach to modeling with an unlimited number of parameters (Gershman & Blei, 2011).

Some statisticians worry that rigorous statistical tools are being displaced by deep learning, which lacks rigor (Friedman, 1998; Matloff, 2014, but see Breiman, 2001; Efron & Hastie, 2016). Assumptions are rarely stated. There are no confidence intervals on the solution. However, performance is typically cross-validated, showing generalization. Deep learning is not convex, but it has been proven that convex networks can compute posterior probability (e.g. Rojas, 1996). Furthermore, machine learning, and statistics seem to be converging to provide a more general perspective on probabilistic inference that combines complexity and rigor.

Some physiologists note that decoding neural activity to recover the stimulus is interesting and useful but falls short of explaining what the neurons do. Some visual psychophysicists note some salient differences between performance of human observers and deep networks on tasks like object recognition and image distortion (Ullman et al. 2016; Berardino et al. 2017). Some cognitive psychologists dismiss deep neural networks as unable to “master some of the basic things that children do, like learning the past tense of a regular verb” (Marcus et al., 1992). Deep learning is slow. To recognize objects in natural images with the recognition accuracy of an adult, a state-of-the-art deep neural network needs five thousand labelled examples per category (Goodfellow et al., 2016). But children and adults need only a hundred labelled letters of an unfamiliar alphabet to reach the same accuracy as fluent native readers (Pelli et al. 2006). Overcoming these challenges may require more than deep learning.

These current limitations drive practitioners to enhance the scope and rigor of deep learning. But bear in mind that some of the best classifiers in computer science were inspired by biological principles (Rosenblatt, 1957; 1958; Rumelhart et al., 1986; LeCun, 1985; LeCun et al. 1989; LeCun et al. 1990; Riesenhuber & Poggio, 1999; and see LeCun, Bengio, Hinton 2015). Some of those classifiers are now so good that they occasionally exceed human performance and might serve as rough models for how biological systems classify (e.g. Yamins, et al. 2014; Khaligh-Razavi & Kriegeskorte, 2014; Ziskind, Hénaff, LeCun, & Pelli, 2014; Testolin, Stoianov, & Zorzi, 2017).

## MILESTONES IN CLASSIFICATION

### Mathematics vs. engineering

The history of machine learning has two threads: mathematics and engineering. In the *mathematical* thread, two statisticians, Fisher and later Vapnik, developed mathematical transformations to uncover categories in data, and proved that they give unique answers. They assumed distributions and proved convergence.

In the *engineering* thread, a loose coalition of psychologists, neuroscientists, and computer scientists (e.g. Turing, Rosenblatt, Minsky, Fukushima, Hinton, Sejnowski, LeCun, Poggio, Bengio) sought to reverse-engineer the brain to build a machine that learns. Their algorithms are typically applied to stimuli with unknown distributions and lack proofs of convergence.

**1936: Linear discriminant analysis**. Fisher (1936) introduced linear discriminant analysis to classify two species of iris flower based on four measurements per flower. When the distribution of the measurements is normal and the covariance matrix between the measurements is known, linear discriminant analysis answers the question: Supposing we use a single-valued function to classify, what linear function *y* = *w*_1_*x*_1_ + *w*_2_*x*_2_ + *w*_3_*x*_3_ + *w*_4_*x*_4_, of four measurements *x*_1_, *x*_2_, *x*_3_, *x*_4_ made on flowers, with free weights *w*_1_, *w*_2_, *w*_3_, *w*_4_, will maximize discrimination of species?^3^ Linear classifiers are great for simple problems for which the category boundary is a hyperplane in a small number of dimensions. However, complex problems like object recognition typically require more complex category boundaries in a large number of dimensions. Furthermore, the distributions of the features are typically unknown and may not be normal.

Cortes & Vapnik (1995) note that the first algorithm for pattern recognition was Fisher’s optimal decision function for classifying vectors from two known distributions. Fisher solved for the optimal classifier in the presence of gaussian noise and known covariance between elements of the vector. When the covariances are equal, this reduces to a linear classifier. The ideal template matcher of signal detection theory is an example of such a linear classifier (Peterson et al., 1954). This fully specified simple problem can be solved analytically. Of course, many important problems are not fully specified. In everyday perceptual tasks, we typically know only a “training” set of samples and labels.

**1953: Machine learning.** The first developments in machine learning were to play chess and checkers. “Could one make a machine to play chess, and to improve its play, game by game, profiting from its experience?” (Turing, 1953). Arthur Samuel (1959) defined *machine learning* as the “Field of study that gives computers the ability to learn without being explicitly programmed.”

**1958: Perceptron.** Inspired by physiologically measured receptive fields, Rosenblatt (1958) showed that a very simple neural network, the perceptron, could learn to classify from training samples. Perceptrons combined several linear classifiers to implement piecewise-linear separating surfaces. The perceptron learns the weights to use in a linear combination of feature-detector outputs. The perceptron transforms the stimulus into a binary feature vector and then applies a linear classifier to the feature vector. The perceptron is piecewise linear and has the ability to learn from training examples without knowing the full distribution of the stimuli. Only the final layer in the perceptron learns.

**1969: Death of the perceptron.** However, it quickly became apparent that the perceptron and other single-layer neural networks cannot learn tasks that are not linearly separable, i.e. cannot solve problems like connectivity (Are all elements connected?) and parity (Is the number of elements odd or even?); people solve these readily (Minsky & Papert, 1969). On this basis, Minsky and Papert announced the death of artificial neural networks.

**1974: Backprop.** The death of the perceptron showed that learning in a one-layer network was too limited. This impasse was broken by the introduction of the backprop algorithm, which allowed learning to propagate through multiple-layer neural networks. The history of backprop is complicated (see Schmidhuber, 2015). The idea of minimization of error through a differentiable multi-stage network was discussed as early as the 1960s (e.g. Bryson, Denham, & Dreyfus, 1963). It was applied to artificial neural networks in the 1970s (e.g. Werbos, 1974). In the 1980s, efficient backprop first gained recognition, and led to a renaissance in the field of artificial neural network research (LeCun, 1985; Rumelhart, Hinton, & Williams, 1986). During the 2000s backprop neural networks fell out of favor, due to four limitations (Vapnik, 1999): **1. *No proof of convergence.*** Backprop uses gradient descent. Gradient descent with a nonconvex cost function with multiple minima is only guaranteed to find a local, not the global minimum of the cost function. This has long been considered a major limitation, but LeCun et al. (2015) claim that it hardly matters in practice in current implementations of deep learning. **2. *Slow***. Convergence to a local minimum can be slow due to the high dimensionality of the weight space. **3. *Poorly specified***. Backprop neural networks had a reputation for being ill-specified, an unconstrained number of units and training examples, and a step size that varied by problem. “Neural networks came to be painted as slow and fussy to train [,] beset by voodoo parameters and simply inferior to other approaches.” (Cox & Dean, 2014). **4. *Not biological***. Lastly, backprop learning may not to be physiological: While there is ample evidence for Hebbian learning (increase of a synapse’s gain in response to correlated activity of the two cells that it connects), such changes are never propagated backwards, beyond the one synapse, to a previous layer. ***5. Inadequate resources***. With hindsight it is clear that backprop in the 80’s was crippled by limited computing power and lack of large labeled datasets.

**1980: Neocognitron,** the first convolutional neural network. Fukushima (1980) proposed and implemented the Neocognitron, a hierarchical, multilayer artificial neural network. It recognized stimulus patterns (deformed numbers) despite small changes in position and shape.

**1987: NETtalk,** the first impressive backprop neural network. Sejnowski et al. (1987) reported the exciting success of NETtalk, a neural network that learned to convert English text to speech: “*The performance of NETtalk has some similarities with observed human performance. (i) The learning follows a power law. (ii) The more words the network learns, the better it is at generalizing and correctly pronouncing new words. (iii) The performance of the networks degrades very slowly as connections in the network are damaged: no single link or processing unit is essential. (iv) Relearning after damage is much faster than learning during the original training…*”

**1989: ConvNets.** Yann LeCun and his colleagues combined convolutional neural networks with backprop to recognize handwritten characters (LeCun et al., 1989; LeCun et al., 1990). This network was commercially deployed by AT&T, and today reads millions of checks a day (LeCun, 1998). Later, adding half-wave rectification and max pooling greatly improved its accuracy in recognizing objects (Jarrett et al., 2009).

**1995: Support Vector Machine (SVM).** Cortes & Vapnik (1995) proposed the support vector network, a learning machine for binary classification problems. SVMs generalize well and are free of mysterious training parameters. Many versions of the SVM are convex (e.g. Lin, 2001).

**2006: Backprop revived.** Hinton & Salakhutdinov (2006) sped up backprop learning by unsupervised pre-training. This helped to revive interest in backprop. In the same year, a supervised backprop-trained convolutional neural network set a new record on the famous MNIST handwritten-digit recognition benchmark (Ranzato et al., 2006).

**2012: Deep learning.** Geoff Hinton says, “It took 17 years to get deep learning right; one year thinking and 16 years of progress in computing, praise be to Intel.” (Cox & Dean, 2014; LeCun, Bengio, & Hinton, 2015). It is not clear who coined the term “deep learning”.^4^ In their book, *Deep Learning Methods and Applications*, Deng & Yu (2014) cite Hinton et al. (2006) and Bengio (2009) as the first to use the term. However, the big debut for deep learning was an influential paper by Krizhevsky et al. (2012) describing AlexNet, a deep convolutional neural network that classified 1.2 million high-resolution images into 1000 different classes, greatly outperforming previous state-of-the-art machine learning and classification algorithms.

## CONTROVERSIES

The field is growing quickly, yet certain topics remain hot. For proponents of deep learning, the ideal network is composed of simple elements and learns everything from the training data. At the other extreme, computer vision scientists argue that we know a lot about how the brain recognizes objects, which we can engineer into the networks before learning (e.g. gain control and normalization). Some engineers look to the brain only to copy strengths of the biological solution, others think there are useful clues in its limitations as well (e.g. crowding).

### Is deep learning the best solution for all visual tasks?

Deep learning is not the only thing in the vision scientist’s toolbox. The complexity of deep learning may be unwarranted for simple problems that are well handled by, e.g. SVM. Try shallow networks first, and, if they fail, go deep.

### Why object recognition?

The visual task of object recognition as has been very useful in vision research because it is an objective task that is easily scored as right or wrong, is essential in daily life, and captures some of the magic of seeing. It is a classic problem with a rich literature. Deep neural nets solve it, albeit with a million parameters. Recognizing objects is a basic life skill, including recognition of words, people, things, and emotions. The concern that the research focus on object recognition might be merely an obsession of the scientists rather than a central task of biological vision is countered by hints that visual perception is biased to interpret the world as consisting of discrete objects even when it isn’t, e.g. when we see animals in the clouds.

Of course, there are many other important visual tasks, including interpolation (e.g. filling in) and extrapolation (e.g. estimating heading). The inverse of categorization is synthesis. Human estimation of one feature, e.g. of brightness or speed, is imprecise and adequately represented by roughly 7 categories (Miller, 1956). For detection of image distortion, a simple model with gain-control normalization is better than current deep networks (Berardino et al. 2017). Scientists, like the brain, use whatever tool works best.

### Deep learning is not convex

A problem is convex if the cost function is convex, i.e. if the line between any two points on the function lies on or above the function. This guarantees that gradient descent will find the global minimum. For some combinations of stimuli, categories, and classifiers, convexity can be proved. In machine learning, kernel methods, including learning by SVMs, have the advantage of easy-to-prove convexity, at the cost of limited generalization. In the 1990s, SVMs were popular because they guaranteed fast convergence even with a large number of training samples (Cortes & Vapnik, 1995). Thus, when the problem is convex, the quality of solution is assured, and one can rate implementations by their demands for size of network and training sample. However, cost functions for deep neural networks are not convex. Unlike convex functions, nonconvex functions can have multiple minima and saddle points. The challenge in high dimensional cost functions is the saddle points, which greatly outnumber the local minima, but there are tricks for not getting stuck at saddle points (Dauphin et al. 2014). Although deep neural networks are not convex, they do fit the training data, and generalize well (LeCun, Bengio, & Hinton, 2015).

### Shallow vs. deep networks

The field’s imagination has focused alternately on shallow and deep networks, beginning with the Perceptron in which only one layer learned, followed by backprop, which allowed multiple layers and cleared the hurdles that doomed the Perceptron. Then SVM, with its single layer, sidelined the multilayer backprop. Today multilayer deep learning reigns; Krizhevsky et al. (2012) attributed the success of AlexNet to its 8-layer depth; it performed worse with fewer layers. Some people claim that deep learning is essential to recognize objects in real world scenes. For example, the “Inception” 22-layer deep learning network won the Image Net Real World Challenge in 2014 (Szegedy et al. 2015).

The need for depth is hard to prove, but, in considering the depth vs. width of a feed-forward neural network, Eldan and Shamir (2016) show that a radial function can be approximated by a 3-layer network with far fewer neurons than the best 2-layer network (also see Telgasky, 2015). Object recognition implies a classification function that assigns one of several discrete values to each image. Mhaskr et al. (2017) suggest that for real-world recognition the classification function is typically compositional, i.e. a hierarchy of functions, one per node, in feed-forward layers, in which the receptive fields of higher layers are ever larger. They argue that scalability and shift invariance in natural images require compositional algorithms. They prove that deep hierarchical networks can approximate compositional functions with the same accuracy as shallow networks but with exponentially fewer training parameters.

### Supervised vs. unsupervised

Learning algorithms for a classifier can be supervised or not, i.e. need labels for training, or don’t. Today most machine learning is *supervised* (LeCun, Bengio, & Hinton, 2015). The images are labeled (e.g. “car” or “face”), or the network receives feedback on each trial from a cost function that assesses how well its answer matches the image’s category. In *unsupervised* learning, no labels are given. The algorithm processes images, typically to minimize error in reconstruction, with no extra information about what is in the (unlabeled) image. A cost function can also reward decorrelation and sparseness (e.g. Olshausen and Field, 1996). This allows learning of image statistics and has been used to train early layers in deep neural networks. Human learning of categorization is sometimes done with explicitly named objects — “Look at the tree!” — but more commonly the feedback is implicit. Consider reaching your hand to raise a glass of water. Contact informs vision. On specific benchmarks, where the task is well-defined and labeled examples are available, supervised learning can excel (e.g. AlexNet), but unsupervised learning may be more useful when few labels are available.

## CURRENT DIRECTIONS

### What does deep learning add to the vision-science toolbox?

Deep learning is more than just a souped-up regression (Marblestone et al., 2016). Like Signal Detection Theory (SDT), it allows us to see more in our behavioral and neural data. In the 1940’s, Norbert Wiener and others developed algorithms to automate and optimize signal detection and classification. A lot of it was engineering. The whole picture changed with the SDT theorems, mainly the proof that the maximum-likelihood receiver is optimal for a wide range of simple tasks (Peterson et al., 1954). In white noise a traditional receptive field computes the likelihood of the presence of a signal matching the receptive field weights. It was exciting to realize that the brain contains 10^11^ likelihood computers. Later work added prior probability, for a Bayesian approach. Tanner & Birdsall (1958) noted that, when figuring out how a biological system does a task, it is very helpful to know the optimal algorithm and to rate observed performance by its *efficiency* relative to the optimum. SDT solved detection and classification mathematically, as maximum likelihood. It was the classification math of the sixties. Machine learning is the classification math of today. Both enable deeper insight into how biological systems classify. Of course, as noted above, SDT is restricted to the case of known signals in additive noise, whereas deep learning can solve real world object recognition like detecting a dog in a photo after training on labeled examples. In the old days we used to compare human and ideal classification performance (Pelli et al. 2006). Today, we also compare human and machine learning. Deep learning is the best model we have today for how complex systems of simple units can recognize objects as well as the brain does. Deep learning, i.e. learning by multi-layered neural networks using backprop, is not just AlexNet but also includes ConvNets and other architectures of trained artificial neural networks. Several labs are currently comparing patterns of activity of particular layers to neural responses in various cortical areas of the mammalian visual brain (Yamins et al. 2014; Khaligh-Razavi & Kriegeskorte, 2014).

### What computer scientists can learn from psychophysics

Computer scientists build classifiers to recognize objects. Vision scientists, including psychologists and neuroscientists, study how people and animals classify in order to understand how the brain works. So, what do computer and vision scientists have to say to each other? Machine learning accepts a set of labelled stimuli to produce a classifier. Much progress has been made in physiology and psychophysics by characterizing how well biological systems can classify stimuli. The psychophysical tools (e.g. threshold and signal detection theory) developed to characterize behavioral classification performance are immediately applicable to characterize classifiers produced by machine learning (e.g. Ziskind, Hénaff, LeCun, & Pelli, 2014; Testolin, Stoianov, & Zorzi, 2017).

### Psychophysics

“Adversarial” examples have been presented as a major flaw in deep neural networks (Mims, 2018; Hutson, 2018). These slightly doctored images of objects are misclassified by a trained network, even though the doctoring has little effect on human observers. The same doctored images are similarly misclassified by several different networks trained with the same stimuli (Szegedy, et al., 2013). Humans too have adversarial examples. Illusions are robust classification errors. The blindspot-filling-in illusion is a dramatic adversarial example in human vision. While viewing with one eye, two finger tips touching in the blindspot are perceived as one long finger. If the image is shifted a bit so that the fingertips emerge from the blindspot the viewer sees two fingers. Neural networks lacking the anatomical blindspot of human vision are hardly affected by the shift (but see Azulay & Weiss, 2018). The existence of adversarial examples is intrinsic to classifiers trained with finite data, whether biological or not. In the absence of information, neural networks interpolate and so do biological brains. Psychophysics, the scientific study of perception, has achieved its greatest advances by studying classification errors (Fechner, 1860). Such errors can reveal “blindspots”. Stimuli that are physically different yet indistinguishable are called *metamers*. The systematic understanding of color metamers revealed the three dimensions of human color vision (Palmer, 1777; Young, 1802; Helmholtz, 1860). In recent work, many classifiers have been trained solely with the objects they are meant to classify, and thus will classify everything as one of those categories, even doctored noise that is very different from all of the images. It is important to train with sample images that represent the entire test set.

## CONCLUSION

Machine learning is here to stay. Deep learning is better than the “neural” networks of the eighties. Machine learning is useful both as a model for perceptual processing, and as a decoder of neural processing, to see what information the neurons are carrying. The large size of the human cortex is a distinctive feature of our species and crucial for learning. It is anatomically homogenous yet solves diverse sensory, motor, and cognitive problems. Key biological details of cortical learning remain obscure, but, even if they ultimately preclude backprop, the performance of current machine learning algorithms is a useful benchmark.

## RESOURCES

We recommend textbooks on deep learning by Goodfellow, Bengio, & Courville (2016) and Ng (2017). There are many packages for optimization and machine learning in MATLAB and Python.

## ACKNOWLEDGEMENTS

Thanks to Yann LeCun for helpful conversations. Thanks to Aenne Brielmann, Augustin Burchell, Kaitie Holman, Katerina Malakhova, Cristina Savin, Laura Suciu, Pascal Wallisch, and Avi Ziskind for helpful comments on the manuscript. We thank our editor, David Brainard, and both reviewers, Nikolaus Kriegeskorte and anonymous, for many constructive suggestions. DGP was supported by NIH grant R01 EY027964.

## GLOSSARY

*Machine learning*: is any computer algorithm that learns how to perform a task directly from examples, without a human providing explicit instructions or rules for how to do so. In one type of machine learning, called “supervised learning,” correctly labeled examples are provided to the learning algorithm, which is then “trained” (i.e. its parameters are adjusted) to be able to perform the task correctly on its own and generalize to unseen examples.
*Deep learning*: is a newly successful and popular version of machine learning that uses backprop (defined below) neural networks with multiple hidden layers. The 2012 success of AlexNet, then the best machine learning network for object recognition, was the tipping point. Deep learning is now ubiquitous in the internet. The idea is to have each layer of processing perform successively more complex computations on the data to give the full “multi-layer” network more expressive power. The drawback is that it is much harder to train multi-layer networks (Goodfellow et al. 2016). Deep learning ranges from discovering the weights of a multilayer network to parameter learning in hierarchical belief networks.
*Neural nets*: are computing systems inspired by biological neural networks that consist of individual neurons learning their connections with other neurons in order to solve tasks by considering examples.
*Supervised learning*: refers to any algorithm that accepts a set of labeled stimuli — a training set — and returns a classifier that can label stimuli similar to those in the training set.
*Unsupervised learning*: discovers structure and redundancy in data without labels. It is less widely used than supervised learning, but of great interest because labeled data are scarce while unlabeled data are plentiful.
*Cost function.*: A function that assigns a real number representing cost to a candidate solution by measuring the difference between the solution and the desired output. Solving by optimization means minimizing cost.
*Gradient descent:*: An algorithm that minimizes cost by incrementally changing the parameters in the direction of steepest descent of the cost function.
*Convexity:*: A real-valued function is called “convex” if the line segment between any two points on the graph of the function lies on or above the graph (Boyd & Vandenberghe, 2004). A problem is convex if its cost function is convex. Convexity guarantees that gradient descent will always find the global minimum.
*Generalization*: is how well a classifier performs on new, unseen examples that it did not see during training.
*Cross validation*: assesses the ability of the network to generalize, from the data that it trained on, to new data.
*Backprop,*: short for “backward propagation of errors”, is widely used to apply gradient-descent learning to multi-layer networks. It uses the chain rule from calculus to iteratively compute the gradient of the cost function for each layer.
*Hebbian learning*: and spike-timing-dependent plasticity (***STDP***). According to Hebb’s rule, the efficiency of a synapse increases after correlated pre- and post-synaptic activity. In other words, neurons that fire together, wire together (Löwel & Singer, 1992).
*Support Vector Machine (SVM)*: is a type of machine learning algorithm for classification. SVMs generalize well. An SVM uses the “kernel trick” to quickly learn to perform a nonlinear classification by finding a boundary in multidimensional space that separates different classes and maximizes the distance of class exemplars to the boundary (Cortes & Vapnik, 1999).
*Convolutional neural networks (ConvNets)*: have their roots in the Neocognitron (Fukushima 1980) and are inspired by the simple and complex cells described by Hubel and Wiesel (1962). ConvNets apply backprop learning to multilayer neural networks based on convolution and pooling (LeCun et al., 1989; LeCun et al., 1990; LeCun et al., 1998).

## Footnotes

1 Admittedly, these networks still demand tweaking of a few parameters, including number of layers and number of units per layer.

2 In the same spirit, “sequential ideal observer” and “accuracy maximization” modeling generalized ideal observer calculations to include a shallow form of supervised learning (Geisler, 1989; Burge & Jaini, 2017).

3 Linear discriminant analysis is an outgrowth of regression which has a much longer history. Regression is the optimal least-squares linear combination of given functions to fit given data and was applied by Legendre (1805) and Gauss (1809) to astronomical data to determine the orbits of the comets and planets around the sun. The estimates come with confidence intervals and the fraction of variance accounted for, which rates the goodness of the explanation.

4 The idea of “deep learning” is not exclusive to machine learning and neural networks (e.g. Dechter, 1986)

